# A causal role for the posterior corpus callosum in bimanual coordination

**DOI:** 10.1101/2025.07.02.662886

**Authors:** Jung Uk Kang, Lawrence H. Snyder, Eric Mooshagian

**Author notes:** Corresponding author: Jung Uk Kang. Department of Behavioral Science, The University of Texas MD Anderson Cancer Center, Houston, TX 77030, USA. Department of Cognitive Science, University of California San Diego, La Jolla, CA 92093, USA.

## Abstract

Inter-areal communication is crucial for brain function. Given the largely contralateral organization of the brain, bimanual coordination likely involves interactions across the two cerebral hemispheres for motor planning and execution. The parietal reach region (PRR) is an early node in the sensorimotor transformation stream. Here we examine the contributions of direct callosal connections between left and right PRR to bimanual coordination. Using manganese-enhanced magnetic resonance imaging, we traced callosal pathways crossing the midline and found that PRR-PRR connections are restricted to the splenium. We then temporarily blocked these fibers with lidocaine while measuring behavioral performance and interhemispheric coherence. Blockade reduced task-specific PRR-PRR coherence during bimanual movements. Behaviorally, blockade sped movement initiation across tasks, consistent with an inhibitory role of interhemispheric communication, reduced the temporal synchrony of bimanual movements to a common target and reduced errors for bimanual movements to separate targets. These findings provide causal evidence that posterior callosal communication supports spatial coordination of bimanual actions but may also constrain independent limb control.

**Significance Statement:** Classic split-brain studies revealed that severing the corpus callosum impairs bimanual coordination, but the specific pathways and mechanisms remain unclear. In macaques, we transiently disrupted the posterior corpus callosum connecting left and right parietal reach regions (PRR), which encode planned contralateral arm movements. This targeted blockade reduced task-specific neural synchrony between PRRs and selectively impaired coordination when both arms reached to a common target, while improving performance when the arms moved to separate targets. Movement initiation was also sped up across tasks, consistent with an inhibitory role of interhemispheric communication. These findings provide causal evidence that posterior callosal communication enables spatially coordinated bimanual movements, extending foundational split-brain insights to defined cortical circuits in a non-human primate model.

## Introduction

Primates excel at planning and coordinating movements (1–4). Tasks that appear to be accomplished using a sequence of independent sensory-to-motor transformations may in fact require advance movement planning and coordination of different body parts (5–9). Scaling a cliff, for example, requires a series of coordinated limb movements to select optimal handholds for ascent. Many steps involve coordinated movements of all four limbs at once, shifting weight and bracing as one reaches to grasp the next hold. Such tasks emphasize the importance of interlimb coordination (10, 11).

Each cerebral hemisphere primarily controls the contralateral limbs (12–17). Given the largely lateralized organization of the brain, bimanual coordination likely requires interactions across the cerebral hemispheres (18–22). Reach planning and execution drives systematic patterns of activity in the functionally defined parietal reach region (PRR) of the posterior parietal cortex (23–27). The population average firing rate in PRR specifically codes planned reaches of the contralateral arm (23, 25, 26), and lesions in PRR impact these movements (14, 28–31). However, beta-band local field potential (LFP) power, reflecting both areal input and local processing (32–34), carries information about both arms (35). Thus, while PRR output primarily encodes contralateral arm information, it receives information about the ipsilateral limb from other areas, perhaps including PRR in the opposite hemisphere (35).

Bimanual tasks provide a means to study how the cerebral hemispheres interact to coordinate the movements of two limbs (36–41). Bilateral recordings made in PRR during bimanual reaching tasks show that interhemispheric LFP-LFP and spike-LFP coherence is modulated by movement type (35). This shared information could arise either from common input to, or from direct communication between left and right PRR.

The corpus callosum is the principal path for information flow between the two cerebral hemispheres (42–44). Gross sectioning of the anterior, posterior, or the whole corpus callosum in humans degrades bimanual coordination (45–48). Yet there is little consensus about how specific callosal pathways support information exchange and complex behaviors (42, 44, 49–51).

Here, we identified and transiently blocked callosal fibers connecting left and right PRR to examine their role in coordinated bimanual behavior. Callosal communication appears to support symmetrical movements to a single target and effectively impedes asymmetrical movements to different targets, consistent with previous clinical studies with callosotomy patients (45–47). Additionally, the modulation of interhemispheric LFP-LFP coherence between left and right PRR based on bimanual movement tasks was significantly reduced during blockade. These results support a causal role of direct interhemispheric communication between left and right PRR in bimanual coordination.

## Results

To establish a baseline, we recorded activity in PRR while animals planned different types of control and coordinated movements (saccades, ipsimanual reaches, contramanual arm reaches, bimanual reaches to the same target, and bimanual reaches to two different targets [Fig. 1]). We identified index cells as single units on the medial bank of the intraparietal sulcus, close to the junction with the parieto-occipital sulcus, that showed strong delay period activity for reaches. We defined PRR as the region spanned by these index cells and classified all task-responsive cells lying within 2 mm of the index cells as PRR cells, regardless of their activity patterns. The population of all such recorded single units shows three levels of delay period activity, similar to previously published results (26). Activity is high when the goal lies in the preferred direction (“preferred”) and when the movement plan includes a reach with the contralateral arm (e.g., a contramanual reach or a coordinated bimanual reach to a single target). Activity is low for any movement away from the preferred direction (“null”). Finally, activity is intermediate when the goal lies in the preferred direction and the movement plan does not include a contramanual reach (i.e., a saccade or ipsimanual reach). Thus, at the population level, PRR neurons distinguish the direction of a planned movement and, for movements in the preferred direction, whether that plan includes a reach with the contramanual arm. In contrast, beta-band LFP power does not distinguish movement direction but does distinguish each of the five movement types described above (Fig. S1A–B; see also Mooshagian et al., 2021) (35). Results from the first animal have been previously reported (Monkey T) (26, 35, 52–54), while results from the second animal replicate that work (Monkey J).

**Fig. 1.**
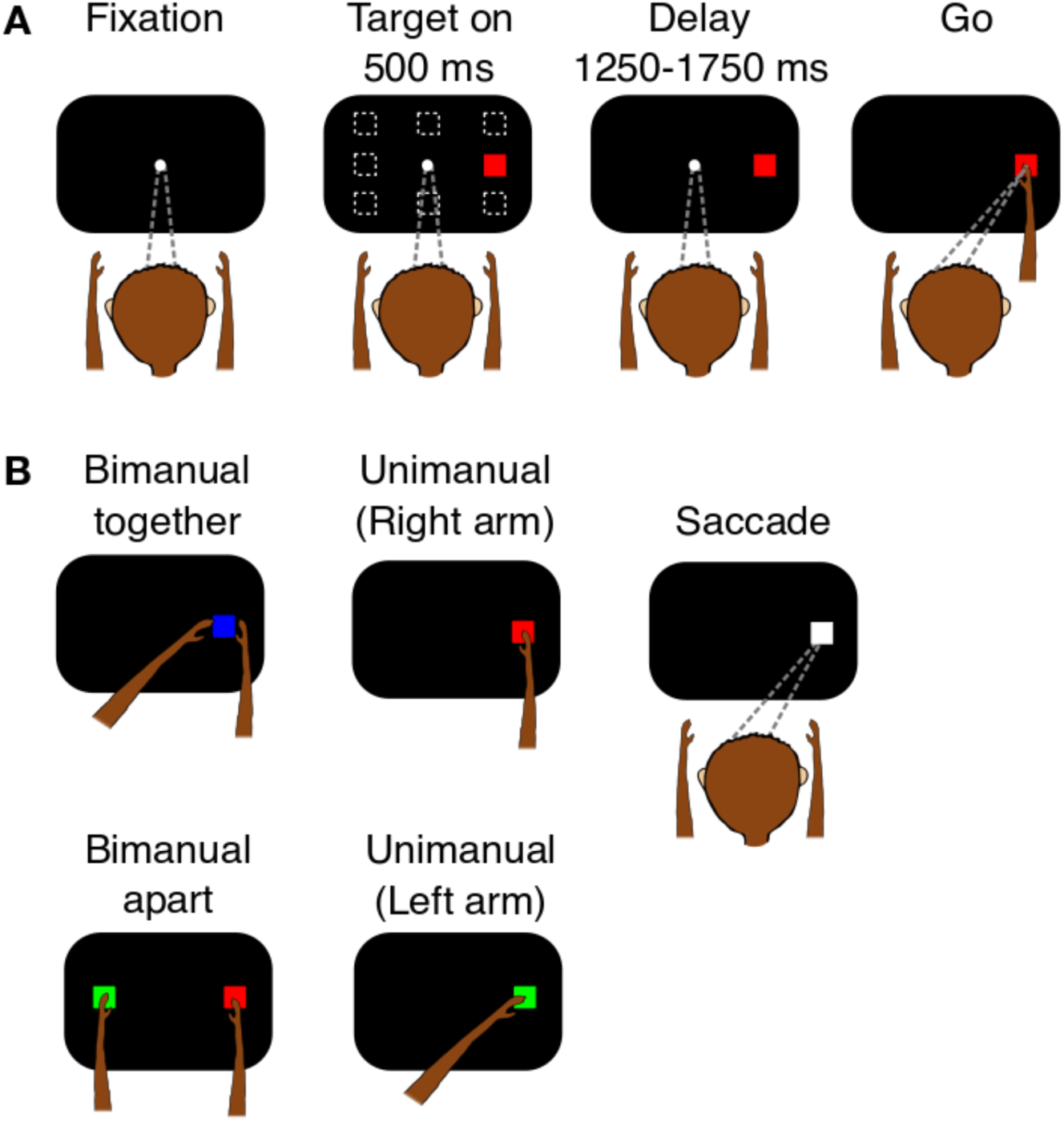
Delayed movement tasks. *(A)* Delayed movement tasks. Animals first fixated on a central target and placed each hand on a home button situated at waist height in front of each shoulder. After an initial fixation period of 500 ms, a peripheral target appeared at one of eight possible peripheral positions (dotted squares). The color and location of the target instructed a particular movement. The target remained visible throughout a variable delay period (1250-1750 ms). After the delay, the disappearance of the fixation target cued the animal to perform the instructed movement. Reproduced from Kang et al. (2024), *Cell Reports*, under CC BY-NC license (54). *(B)* Types of movement tasks. Animals performed two types of bimanual movements: “Bimanual-together” (two arms, one target) and “Bimanual-apart” (two arms, two targets). Bimanual-together movements were both spatially and temporally coordinated, that is, directed towards the same target, whereas bimanual-apart movements were not spatially coordinated (different targets). Unimanual reaches and saccade-only trials served as controls.

The action potentials from single units are directional but LFP is not. PRR, unlike many other posterior areas, contains roughly equal numbers of intermingled cells with contralateral or ipsilateral response fields (RFs) (26, 55). An LFP recording combines (dendritic, somatic, and axonal) signals from many cells, and therefore directional specificity is lost. However, the fact that LFP retains sensitivity to the five types of movements is not so easily explained. Mooshagian and colleagues proposed that each PRR codes reach plans for just the contralateral arm (35). Cells from the opposite PRR provide information about what the other arm (the ipsilateral arm) is doing, allowing PRR to adjust the contralateral arm plans, thereby providing a neuronal mechanism for bimanual coordination. A simple model shows that this can explain the levels of LFP activity seen in PRR (Supplementary Fig. 13 of Mooshagian et al., 2021) (35).

To examine the role of callosal connections between left and right PRR in bimanual coordination, we recorded spikes and LFPs in both hemispheres simultaneously and compared reaching behavior and neuronal activity before and during blockade of homotopic callosal connections between left and right PRR. To identify the relevant pathways, we injected manganese into PRR and then used *in vivo* manganese-enhanced magnetic resonance imaging (Mn-MRI) to trace white-matter pathways between PRR in each hemisphere (56, 57). Axons from PRR crossing to the opposite hemisphere in the posterior portion of the brain were restricted to the splenium (Fig. 2A and Fig. S2A). We reversibly blocked these callosal pathways by injecting 5 μL of a 2% lidocaine solution into the splenium at a rate of 0.15 μL/minute. We confirmed our targeting by co-injecting manganese with the lidocaine and imaging the animal to confirm that we had injected the same area that we had identified in the tract-tracing experiments (Fig. S2B–C).

**Fig. 2.**
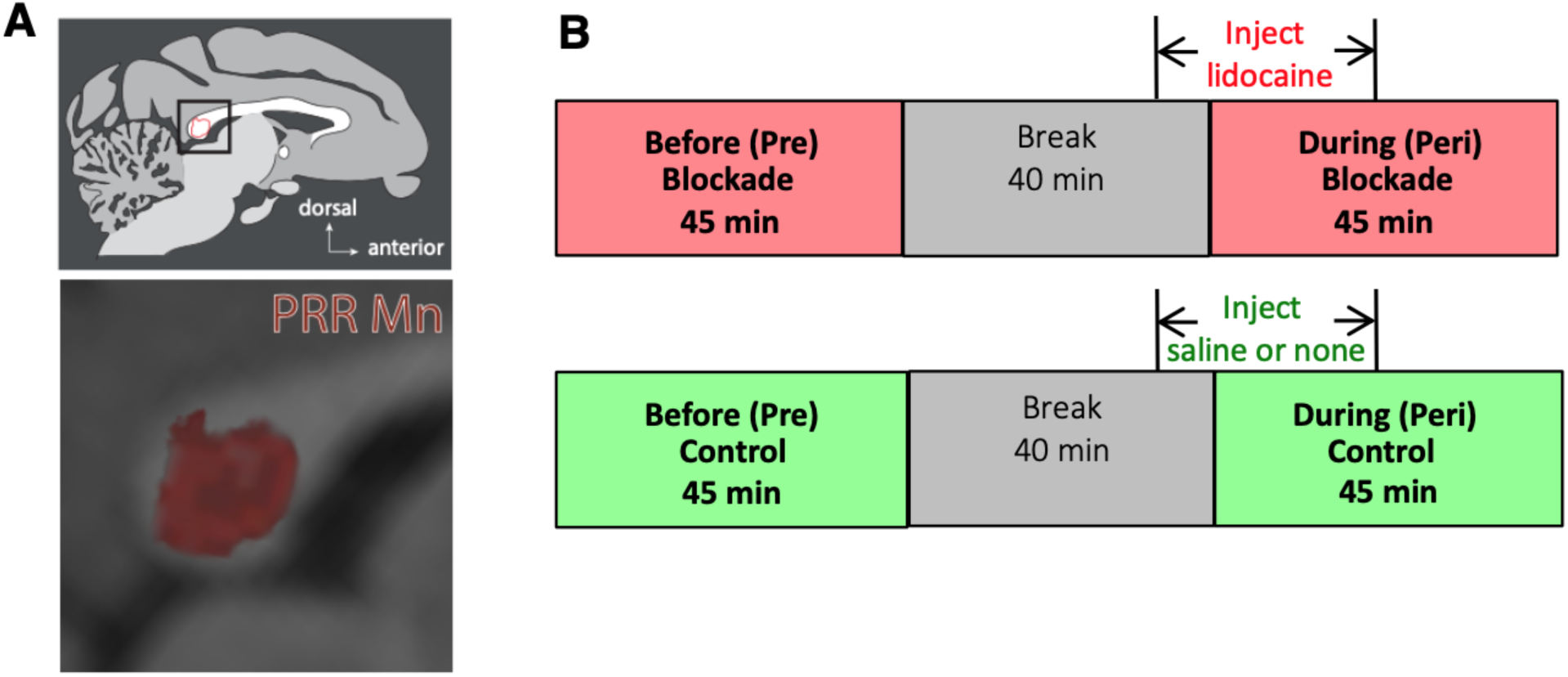
Experimental design. *(A)* Localization of callosal fibers connecting left and right PRR using Mn-MRI. Top, callosal fibers projecting from right PRR and through the corpus callosum were restricted to the splenium (red outline). Bottom, an expanded view. Full Mn-MRI scan showing the anterograde transport of Mn from PRR is in Fig. S2. *(B)* Timeline of experimental and control sessions. Each session comprised two blocks of 400 or 480 trials each. One block was completed before (Pre Blockade) and one during (Peri Blockade) the injection with a 40-minute break between the two. Experimental and control sessions differed by whether a real or sham injection was conducted. In experimental sessions, lidocaine was injected (5.0 μL for Monkey J and 5.5 μL for Monkey T of 2% lidocaine solution; rate = 0.15 μL/minute). In control sessions, either saline or nothing was injected. Peri Blockade blocks began once 1.0 μL of lidocaine was injected (6.6 minutes) and ended 15 minutes after the injection was completed (see Methods).

We obtained two blocks of 400-480 interleaved trials of the five delayed movement tasks (Fig. 2B). One block (Pre) was collected before the injection and a second block (Peri) began once 1.0 μL of lidocaine was injected and ended within 20 minutes after the end of the injection. We also ran control sessions in which a guide tube was lowered and either saline or nothing was injected. We compared behavioral performance and neural activity between callosal blockade and control sessions.

### Callosal pathways facilitate temporal synchronization and spatial coordination of bimanual movements

We first tested whether posterior callosal pathways might help synchronize movements of the two arms. We quantified bimanual synchrony by measuring the absolute difference in reaction times (RTs) between the two arms, with smaller differences indicating greater synchrony. Fig. 3A shows how synchrony changes with callosal blockade for the two bimanual movement tasks. Upward (positive-going) bars indicate an increase in synchrony. During callosal blockade, bimanual-together movements became less synchronous by −7.1 ± 3.6 ms (pooled t-test, *p* = 0.054, both animals; −7.3 ms ± 4.3 ms, Monkey J and −5.1 ms ± 5.2 ms, Monkey T; Fig. S3A and Fig. S3D). Critically, blockade had no effect on the synchrony of bimanual apart movements (−0.7 ± 3.0 ms; pooled t-test, *p* = 0.824). A non-specific effect of blockade on movement timing would degrade the synchrony of both types of movements. These results are therefore consistent with callosal pathways helping to synchronize spatially coordinated movements of the two arms to a single target.

**Fig. 3.**
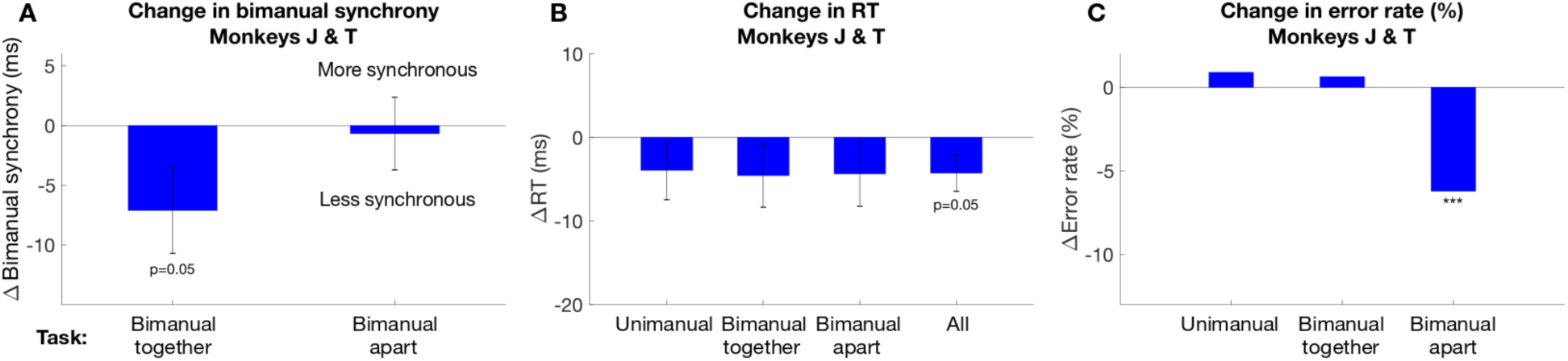
Behavioral performance in callosal blockade and control sessions. *(A)* Change in bimanual synchrony as a function of bimanual movement type and session. We assayed the effect of callosal blockade on the absolute reaction time difference between the two arms, Δ(|RT_Left_ – RT_Right_|) in Peri minus Pre blocks. We compared the change in bimanual synchrony from Pre to Peri blocks in control (*n* = 24) and blockade (*n* = 22) sessions. Positive-going values indicate higher bimanual synchrony during callosal blockade than in control sessions. In bimanual-together movements, we observed significantly less synchronous bimanual movements in blockade sessions compared to control sessions (*p* = 0.05, pooled t-test). No significant changes were observed in bimanual-apart trials. *(B)* Change in RT. Callosal blockade sped movement initiation (shorter RTs) compared to control for all three movement types. This speeding effect was statistically significant when averaged across all movement types (*p* = 0.05, pooled t-test). ΔRT (Peri - Pre) is calculated as the difference between the median RT in the Peri block and the median RT in the Pre block for each movement type (Unimanual, Bimanual-together, Bimanual-apart)in control versus callosal blockade sessions. Error bars in *(A)* and *(B)* denote the standard error of the mean (SEM). Changes in bimanual synchrony and RT in control and blockade sessions are shown separately in Fig. S4. *(C)* Change in error rate. Callosal blockade improved bimanual-apart movements compared to control (*p* < 0.001, logistic regression). ΔError rate (Peri - Pre) is calculated as the difference between error rates in the Peri block and Pre block. Error rate in each block is calculated from data concatenated across all sessions. We compared the change in error rates in control and blockade sessions. We performed logistic regression with block (Pre, Peri) and condition (Control, Blockade) as two factors, and show the statistical significance of the interaction term of the two factors. *** *p* < 0.001

Next, we considered the mechanism by which callosal communication helps synchronize the two arms. We hypothesized that callosal pathways either slow down the faster arm or speed up the slower arm to achieve temporal coordination. Compared to control sessions, callosal blockade resulted in faster RTs across all tasks, including unimanual reaches (Fig. 3B). When pooled across tasks, the effect was significant (−4.3 ms ± 2.2 ms, pooled t-test, *p* = 0.05). This effect was present in both animals, but reached significance only in Monkey J (−7.8 ms ± 3.2 ms, *p* = 0.016, pooled t-test; Fig. S3B). In Monkey T, the effect trended towards faster RTs in a reaction time task in which the animal could move as soon as the target appeared (−9.0 ± 6.1 ms, *p* = 0.149, pooled t-test; *n*=10 blockade sessions, *n*=10 control sessions; only Monkey T performed this task). Thus, blocking callosal transmission speeds up reaching movements. For spatially coordinated movements (bimanual-together), RTs became faster but less synchronous. For spatially distinct movements (bimanual-apart), RTs also became faster but without significant changes in synchrony. These results indicate that callosal communication supports bimanual temporal coordination by slowing down the faster arm rather than speeding up the slower arm.

We also assessed effects of blockade on spatial accuracy. We used capacitive proximity sensors rather than a touch screen or similar device to sense multiple independent touches. As a result, we do not have precise estimates of spatial endpoints. However, we know from monitoring reaches on real time video that almost all arm-related errors occurring after the go cue resulted from the animal landing off-center on the correct capacitive switch, and not because the wrong button was targeted (see Methods and Table S1 for details). As a result, the rate of arm errors occurring after the go cue provides a proxy for reach accuracy. Blockade improved accuracy for bimanual-apart movements (*p* < 0.001, logistic regression) and did not affect unimanual and bimanual-together movements. When separated by animal, the bimanual-apart effect was significant in Monkey T, who also showed a significant improvement in bimanual-together movements (*p* < 0.01, logistic regression; Fig. S3F). Monkey J showed non-significant improvements in both types of bimanual movements (Fig. S3C). Taken together, the effects on synchrony and spatial accuracy suggest that callosal communication helps yoke the two arms together and interferes with bimanual movements to different targets, such that blocking the callosum impairs bimanual-together movements and improves bimanual-apart movements. This is reminiscent of results from patients with posterior callosotomy, in whom symmetric arm movements are impaired while asymmetric movements are improved (45, 47).

We performed similar analyses on synchrony, RT, and error rates as a function of laterality and symmetry (see Methods and Fig. S5). Movements were categorized as crossed or uncrossed based on the neuroanatomical relationship between the stimulus and movement. In crossed movements, each arm reached to a target in the contralateral visual hemifield, meaning that the hemisphere that processed the spatial location and one that controls the movement were different, requiring interhemispheric transfer. In uncrossed movements, the stimulus and motor planning the stimulus and the motor planning occurred in the same hemisphere, and therefore did not require interhemispheric transfer. Movements were categorized as symmetric or asymmetric based on whether the arms moved in a spatially mirror-symmetric fashion (e.g., toward mirror opposed targets) or toward the same lateralized target. Compared to control sessions, callosal blockade sessions resulted in reduced synchrony during asymmetric movements (Fig. S6), no significant change in RT (Fig. S7), and fewer errors in crossed movements (Fig. S8). The observed reduction in synchrony for asymmetric movements is consistent with the notion that callosal communication facilitates temporal synchronization of bimanual asymmetric movements to a single target in either visual hemifield. Fewer errors in crossed bimanual movements during callosal blockade indicate that intact callosal communication may be disruptive for bimanual coordination when each arm moves to a target in the contralateral visual hemifield.

### Callosal blockade greatly reduces task-specific modulation of interhemispheric LFP-LFP coherence

Our behavioral results indicate that communication between the left and right PRR helps support bimanual coordination. We looked for neural correlates of this communication by measuring the effect of callosal blockade on interhemispheric LFP-LFP coherence (see Methods) (58). Beta-band LFP activity often correlates with motor control variables (34, 59–65), and interhemispheric LFP-LFP coherence in this band shows bimanual task-specific modulation (22, 35). Interhemispheric coherence is significantly stronger when preparing bimanual-together than bimanual-apart movements and is intermediate when preparing unimanual movements or saccades. This pattern suggests that interhemispheric communication increases when both arms move toward the same target and decreases when each arm moves to a different target. Alternatively, common input may increase for same-target movements and decrease for movements to different targets. If interhemispheric communication is at play, and if it occurs via direct connections between the left and right PRR, then blocking callosal pathways should disrupt or abolish task-specific modulation of LFP coherence, consistent with the effect of blockade on behavior.

Interhemispheric coherence is shown in Fig. 4 and Fig. S9. Coherence was tested at 2 Hz intervals from 20 to 100 Hz, with a focus on the 20-30 Hz frequency band found to be of particular interest in our previous studies (35, 54). Consistent with previous results, baseline coherence was significantly stronger while planning a bimanual-together movement compared to a bimanual-apart movement at each individual frequency from 20 to 32 Hz (*p* < 0.01) as well as in the *a priori* 20-30 Hz band (difference of 0.046 ± 0.007, *p* < 0.001; Fig. 4). Callosal blockade reduced this difference by 70% (*p* < 0.001) though it did not completely disappear, particularly at the lowest frequencies tested (0.014 ± 0.006, *p* < 0.02).

**Fig. 4.**
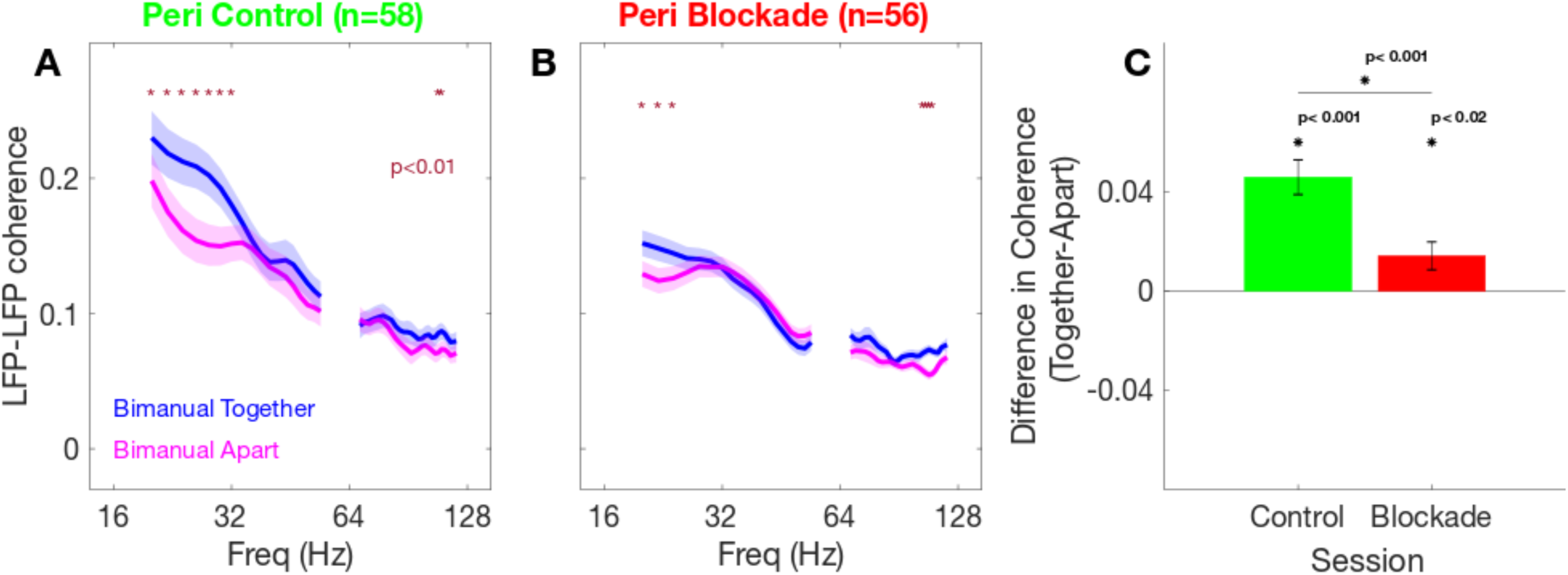
LFP-LFP coherence between left and right PRR in control and callosal blockade sessions. *(A)* Coherence is higher for bimanual-together movements than for bimanual-apart movements at 20-32 Hz. Red asterisks denote differences of *p* < 0.01, tested at 2 Hz intervals averaged over 58 pairs of electrode contacts in each hemisphere. Shaded regions represent ±SEM. *(B)* During the Peri callosal blockade blocks, the bimanual-task specific difference in coherence is reduced at low frequencies and abolished at higher frequencies. Format as in a. *(C)* Blocking the corpus callosum significantly reduced the task-dependent difference in coherence measured from 20-30 Hz (*p* < 0.001, pooled t-test). This frequency range, 20-30 Hz, was specified prior to the start of data collection to avoid multiple comparison issues. Error bars are ±SEM.

We initially hypothesized that callosal communication can explain why, despite single unit activity only distinguishing movements that include the contralateral arm, beta-band LFP modulations are distinct for each of the five movement types that we tested. This hypothesis is supported by the fact that, after callosal blockade, interhemispheric coherence is no longer strongly modulated by whether the two arms move together or apart. However, the beta-band LFP activity continues to show distinct modulations for different patterns of arm movements (Fig. S1C–D).

We performed a similar analysis using data from the left and right lateral intraparietal area (LIP) from one animal. The lidocaine injections were targeted based on tract-tracing from PRR, but it is likely that callosal fibers from LIP were also affected. LIP shows no task-specificity in coherence at baseline and no evidence for a change with callosal blockade (Fig. S10).

## Discussion

Effective bimanual coordination depends on interhemispheric communication via the corpus callosum. Here, we examined the contributions of callosal connections between left and right PRR in mediating these effects. To identify the specific callosal channels involved, we first mapped interhemispheric projections using manganese-enhanced MRI tract-tracing. This revealed that PRR-PRR callosal projections are restricted to the splenium. We then reversibly blocked these splenial fibers and assessed behavioral and neural consequences. Compared to baseline, callosal blockade reduced the temporal synchrony of bimanual-together movements (Fig. 3A), sped up movement initiation across all tasks (Fig. 3B), and reduced errors in bimanual-apart movements (Fig. 3C), suggesting an inhibitory role for callosal communication (44, 66, 67). Neural recordings were complementary to the behavioral findings: task-specific LFP-LFP coherence between left and right PRR was selectively reduced during bimanual tasks during callosal blockade (Fig. 4). Together, these results suggest that the posterior corpus callosum actively modulates bimanual movement planning.

The corpus callosum is the largest white-matter commissure, integrating sensory, motor, and cognitive processes across the cerebral hemispheres. It is topographically organized, with distinct regions supporting specific interhemispheric functions (68–71). Studies of split-brain patients who have undergone callosotomy surgeries provide valuable insights into hemispheric interactions. Studies of these patients implicate the corpus callosum in bimanual coordination (47, 48, 72–74) with deficits emerging in novel or complex tasks while well-learned movements remain intact (48, 75). However, prior studies have either assessed behavior without measuring neural changes, or examined interhemispheric connectivity without linking it to behavior (76–82). This study directly establishes a causal relationship between bimanual coordination and interhemispheric connectivity.

Complete commissurotomy limits the ability to determine specific cortical contributions to interhemispheric interactions. Studies of partial or staged commissurotomy have revealed region-specific contribution of the corpus callosum to bimanual coordination. Severing the anterior corpus callosum primarily affects temporal synchronization and movement initiation (45, 46, 83, 84). In contrast, severing the posterior corpus callosum impairs spatial coordination and directional control (45, 47, 85). Such approaches offer only coarse localization and are irreversible, limiting their utility in pinpointing contributions of specific callosal pathways. By focally and reversibly blocking the posterior corpus callosum, we specifically evaluated the role of callosal pathways between the two PRRs in neural communication and behavioral performance.

Our results indicate that the posterior corpus callosum facilitates temporal coordination of spatially coordinated bimanual movements. Blocking the splenium disrupted synchrony in bimanual-together movements but did not affect synchrony in bimanual-apart movements, suggesting its role is specific to spatially coordinated actions rather than independent limb control. This aligns with prior studies implicating the posterior corpus callosum in the spatial coordination of bimanual movements (45, 48, 85).

Callosal blockade also led to faster movement initiation across all tasks, suggesting that interhemispheric communication, specifically between homotopic areas such as PRR, normally exerts a functionally inhibitory influence on movement onset. Interestingly, error rates decreased selectively during bimanual-apart movements during blockade. This finding is consistent with case studies of posterior callosotomy patients, where asymmetric or independent limb movements sometimes improve (45, 47).

Motor behavior appears to begin from a “mirror” program: during movement planning, regions such as dorsal premotor cortex and supplementary motor area exhibit bilateral activation (86–88). To produce lateralized, unimanual actions, the brain engages interhemispheric inhibitory mechanisms, whereby the active motor cortex suppresses its counterpart (ipsilateral to the intended movement), typically via excitatory callosal projections that engage inhibitory interneurons in the opposite hemisphere (89–91). However, the corpus callosum is not the only determinant of motor lateralization. Congenital absence of the corpus callosum can produce a syndrome in which movements on both sides of the body become coupled, suggesting that additional anatomical pathways contribute to the ability to produce lateralized actions (92). These likely include subcortical structures such as the corticospinal tract and the spinal cord (93–95), but their roles remain to be fully defined.

While our tasks were not specifically designed to assess bimanual movement symmetry, we were able to classify movement types post hoc based on both symmetry and laterality. Callosal blockade had no significant effect on movement errors or RTs when movements were grouped by symmetry (Fig. S7 and Fig. S8). However, we observed a significant effect of blockade when bimanual movements were grouped by laterality, specifically whether they were crossed (Fig. S7). Because each cerebral hemisphere primarily processes spatial information from the contralateral visual hemifield, crossed movements typically require interhemispheric transfer since the hemisphere that perceives the spatial target must communicate with the opposite hemisphere to initiate the motor response. Thus, one might expect callosal blockade to impair performance of crossed movements. Instead, we found that callosal blockade significantly reduced errors in crossed bimanual movements. This improvement suggests that interhemispheric communication may introduce interference in this condition, and that reducing callosal input may improve performance by allowing each hemisphere to process and respond independently. These findings also imply that other cortical or subcortical pathways may be sufficient to support lateralized action planning.

Task-specific LFP-LFP coherence between left and right PRR was reduced or abolished during callosal blockade, indicating a loss of interhemispheric communication (Fig. 4 and Fig. S9). This disruption was specific to bimanual tasks and suggests that interhemispheric coupling of left and right PRR contributes to coordinating bilateral movement plans. The loss of beta-band task-specific coherence corresponds with the decrease in temporal synchrony for bimanual-together movement RT, suggesting that interhemispheric communication plays a role in maintaining coordinated timing between the arms.

Mooshagian and colleagues found that PRR unit activity primarily reflects contralateral arm movement plans, whereas local beta-band LFP encodes both contralateral and ipsilateral arm movements (35). A key hypothesis was that ipsilateral arm information in PRR beta-band LFP activity originates from PRR in the opposite hemisphere via callosal projections, an idea supported by the interhemispheric PRR-PRR beta coherence findings. However, splenium blockade did not abolish local beta-band task specificity in PRR (Fig. S1C–D). This suggests that beta-band activity in PRR does not rely exclusively on direct callosal input from the opposite PRR.

One explanation is that callosal blockade is incomplete, leaving many alternative interhemispheric communication pathways intact. The splenium itself may not be completely blocked by our method, and other callosal pathways or subcortical connections may still contribute to interhemispheric movement encoding. Mooshagian and colleagues (35) (see their Supplementary Table 3) proposed that beta power differentiation is due to a mixture of local and contralateral input, with even a small amount (5-20%) of contralateral input sufficient to maintain movement type selectivity in beta-band activity. The persistence of PRR beta-band LFP selectivity following splenium blockade is therefore consistent with the idea that a small amount of contralateral arm information – whether via residual splenial fibers, alternative callosal channels, or extra-callosal pathways – is sufficient to maintain beta-band movement encoding. An alternative explanation is that the beta-band LFP task selectivity reflects differences in the visual input. This could explain the difference between tasks with one versus two targets, but not the differences between single target tasks.

Theories of callosal function propose that interhemispheric communication can be either excitatory or inhibitory, with human studies providing evidence of both mechanisms (35, 53, 96–98). Faster movement initiation across tasks suggests that callosal projections may normally exert an inhibitory influence on movement initiation. Conversely, the disruption of synchrony in bimanual-together movements suggests that PRR-PRR callosal projections facilitate temporal coordination in spatially aligned movements, possibly through excitatory coupling. The fact that bimanual-apart movements were not affected suggests that this coordination is specific to movement requiring spatial and temporal alignment, and is not a general property of bimanual coordination. It is important to keep in mind that the functional effects of interhemispheric communication do not constrain whether the callosal fibers themselves are excitatory or inhibitory (44, 50).

A limitation of our approach is that callosal blockade likely affects more than just PRR-PRR connections, potentially impacting nearby areas such as LIP (99, 100). A preliminary study using the same methods indicates substantial overlap between callosal pathways from LIP and PRR within the splenium. While our focal blockade altered eye-hand coordination behaviorally, it did not affect neural communication between PRR and LIP or between LIP and LIP in each hemisphere (Fig. S10; J. Kang, E. Mooshagian, L. H. Snyder. Temporal eye-hand coordination may be subserved by connections between PRR and LIP. Neuroscience Meeting Planner. Washington, DC: Society for Neuroscience, 2023. Online.). More precise targeting methods, such as optogenetics or chemogenetics, could clarify the specific role of PRR-PRR callosal pathways in motor coordination (101, 102).

Understanding long-range communication between different brain regions is essential for both basic neuroscience and clinical applications (103–106). Our Mn-MRI approach for mapping specific callosal pathways can be extended to investigate interhemispheric working memory transfer, white-matter contributions to visual processing, or other forms of hemispheric integration (107–109). Future studies using varied behavioral paradigms and selective circuit manipulations will be necessary to further elucidate the neural and behavioral contributions of long-range communication.

## Materials and Methods

### Experimental model and subject details

All procedures conformed to the Guide for the Care and Use of Laboratory Animals and were approved by the Washington University Institutional Animal Care and Use Committee. Two male rhesus macaques (Macaca mulatta), Monkey J (7-year-old male, 14.5 kg) and Monkey T (20-year-old male, 9.0 kg), were used in the study.

### Apparatus

Head-fixed animals sat in a custom-designed monkey chair (Crist Instrument, Hagerstown, MD) with an open front for unimpaired reaching movements with both arms. Visual stimuli were back-projected by an LCD projector onto a translucent plexiglass screen mounted vertically ∼40 cm in front of the animal. The room was otherwise dark. Eight target positions were organized in a rectangle centered on the fixation point, each target ∼8 cm (11°) or ∼11 cm (15°) from the center fixation point. A small piece of plexiglass (5 cm × 1 cm) oriented in the sagittal plane was mounted on the front of the projection screen to bisect the touching surface at each target location. The animals were trained to reach with the left hand to the left side of the divider and the right hand to the right side. Touches were monitored every 2 ms using 16 capacitive sensors, mounted on the back of the screen, one sensor on each side of each of the 8 possible target locations to sense reach endpoints. Thus, each hand activated a unique capacitive sensor, even when both hands reached the same target. Two additional sensors, one sensor at each of the two home pads, were used to monitor the animals’ reach starting positions and one sensor on each side of each of the 8 possible target locations to sense reach endpoints. Thus, each hand activated a unique capacitive sensor, even when both hands reached the same target. Eye position was monitored using the 120 Hz ISCAN eye-tracking laboratory (ETL-400). Animals were monitored in the testing room using two infrared cameras equipped with infrared illuminators, aimed at the plexiglass screen and the home pads.

### Behavioral tasks

The task design and the movement conditions are illustrated in Fig. 1. The animals performed delayed saccade-only movements or coordinated eye plus arm movements using the left, right, or both arms. Initially, the animals fixated on a circular white stimulus (1.5° x 1.5°) centered on the screen. Their left and right hands touched “home” pads situated at waist height, 20 cm in front of each shoulder. After maintaining fixation (± 5°) and initial arm positions for 500 ms, one or two peripheral targets (5° × 5°) appeared on the screen for 1250-1750 ms, during which fixation was required. Next, the fixation target shrank to a single pixel, cueing the animal to move to the peripheral target(s) in accordance with target color and a previously trained code. A green (red) target instructed a reach with the left (right) arm. A red target instructed a reach with the right arm. A blue target instructed a combined reach with both arms (“bimanual-together”). The simultaneous appearance of red and green targets cued a “bimanual-apart” reach, where each arm reached its respective target. For bimanual-apart reaches, the two targets appeared at diametrically opposite locations across the central fixation and the arms could therefore be crossed or uncrossed. A white target instructed a saccade without a reach. To ensure natural coordination, animals were never trained to make arm movements without moving the eyes. All single-target reach trials required an accompanying saccade to the target, while saccades were optional (but almost always performed) for two-target reach trials.

All trial types were randomly interleaved within sets of 40 trials (one each per condition [5 task types] and [8 target configurations]). During saccade and unimanual reach trials, the non-moving hand(s) remained on the home button(s). In bimanual together trials, the left and right hands were required to hit their target within 275-325 ms (Monkey J; Bimanual together), 225 ms (Monkey T; Bimanual together), 300-500ms (Monkey J; Bimanual apart), or 300 ms (Monkey T; Bimanual apart). Animals were required to maintain their hand(s) on the final target(s) for 300 ms. The spatial tolerance for saccades was ±5°. If an error occurred (a failure to achieve or maintain the required eye or hand positions), the trial was aborted, and short time-out ensued (1500ms for early fixation breaks and 500ms for targeting errors). Successful trials were rewarded with a drop of water or juice.

### Manganese-enhanced magnetic resonance imaging (Mn-MRI)

Callosal pathways connecting left and right PRR were identified in one animal (Monkey T) with *in vivo* Mn-MRI (56, 57, 110). Two 33-gauge cannulae attached to 10-μL Hamilton syringes were lowered to two locations in the right PRR. Next, 1.0 μL of 0.3M MnCl_2_(H_2_O)_4_ in 0.9% sterile saline solution buffered with 10 mM Tris-HCl (pH ∼7.4) was injected into each site using a microinjection pump (Harvard Apparatus). We acquired MRI images 24- and 48-hours after the injection. A subtraction image (48 hr - 24 hr) showed anterograde transport of Mn^2+^ from right PRR. Callosal pathways from right PRR to the left hemisphere were restricted to the splenium (Fig. S2A).

### Reversible callosal blockade

We considered the onset time, duration of inhibitory effects of 2% lidocaine, and the injection volume necessary to observe behavior and neural effects, in designing the Peri block. Pilot experiments established that 5.0 μL for Monkey J and 5.5 μL for Monkey T were effective volumes for observing changes in behavior and neural activity. These effects persisted throughout the blockade experiment.

The effects from lidocaine injection have faster onset and shorter duration than muscimol (GABA-A receptor agonist) (14, 28, 110, 111). Neural activity is effectively silenced 2 minutes after lidocaine injection, and neurons regain their initial activity 30 minutes after lidocaine injection (112, 113). To ensure blockade was in effect, the Peri block commenced only after 1.0 uL of lidocaine had been injected (6.6 minutes). To maintain blockade at a constant level throughout the block, lidocaine was continuously injected at a constant rate until a total volume of 5.0 μL (Monkey J) and 5.5 μL (Monkey T) was reached. Peri blockade blocks were completed within 20 minutes after lidocaine injection.

Each blockade session proceeded as follows. A 33-gauge injection cannula attached to a 10-μL Hamilton syringe was lowered to the splenium. At the same time, recording electrodes were lowered to left and right PRR. The electrodes and cannula were allowed to settle for 15 minutes. Next, animals completed a block of 400 trials (Monkey J) or 480 trials (Monkey T) (‘Pre’, 45 minutes). Next, there was a 40 minute break. Following the break, the Peri block began. The details of the Peri block were informed by the onset time, duration and the injection volume necessary to observe behavior and neural effects. Seven minutes before the Peri block commenced, we began injecting 2% lidocaine at a rate of 0.15 μL/min using a microinjection pump (Harvard Apparatus). Next, animals completed a second block of 400 trials (Monkey J) or 480 trials (Monkey T) (‘Peri’, 45 minutes). The Peri block began once 1.0 μL of 2% lidocaine was injected (6.6 minutes). A total of 5.0 μL (Monkey J) or 5.5 μL (Monkey T) of lidocaine was injected, ending 27 (Monkey J) or 30 minutes (Monkey T) into the Peri block.

Blockade and control sessions differed only by whether lidocaine or a sham injection was performed. For control sessions, either 0.9% saline solution was injected (2 sessions for each Monkey) or nothing (sham) was injected (13 sessions for Monkey J; 7 sessions for Monkey T). In the sham sessions, the procedure was identical to a control session with 0.9% saline solution except that the injection apparatus was set up without an injection cannula.

In a subset of sessions, the experimenter was blinded to the session type (blockade or control). For Monkey J, we conducted 1 blinded blockade and 1 blinded control session out of 12 blockade and 15 control sessions, respectively. For Monkey T, we conducted 2 blinded blockade and 2 blinded control sessions out of 10 blockade and 9 control sessions, respectively.

### Injection localization with MRI

We administered manganese injections to confirm our focal blockade coordinates. For Monkey J, we injected 0.08 μL of 0.08M MnCl_2_(H_2_O)_4_ in 0.9% sterile saline solution. For Monkey T, we used 2 μL of 0.06M MnCl_2_(H_2_O)_4_ in 2% lidocaine solution. Both solutions were buffered with 10 mM Tris-HCl (pH ∼7.4). We acquired structural MRI images within 6 hours after the injection as specified in the section *Mn-MRI*. We observed isolated halos in the splenium in both animals and confirmed the accuracy of our injection sites (Fig. S2B–C).

### Electrophysiological recordings

Simultaneous recordings were made from the left and right hemispheres of Monkeys J and T. Each animal had two recording chambers centered approximately at 8 mm posterior to the ear canals and 12 mm lateral of the midline on each side, placed flush to the skull. Anatomical magnetic resonance images were used to localize the medial bank of the intraparietal sulcus. Extracellular recordings were made using two glass-coated tungsten electrodes (Alpha Omega, Alpharetta, GA; impedance 0.5-3.0 MΩ at 1kHz) for both monkeys, or a 32-channel multi-contact electrode (NeuroNexus, Ann Arbor, MI; contacts 200 μm apart; impedance ∼1.25 MΩ at 1kHz) for Monkey J, inserted through a steel guide tube into PRR in each hemisphere. Neural signals from the four single-contact electrodes or the two multi-contact electrodes were processed and saved using the Plexon MAP system (Plexon, Inc.) for Monkey T and the Open Ephys acquisition system (114) for Monkey J. In the Plexon MAP system, signals were passed through a pre-amplifier and then separated into two signal paths for LFP and spikes. The LFP channel was band-pass filtered between 0.7 and 300 Hz and digitized at 1 kHz. The spike channel was band-pass filtered between 100 Hz and 8 kHz and digitized at 25 kHz. In the Open Ephys system, signals were passed through a pre-amplifier and band-pass filtered between 2.5 and 7603.8 Hz and digitized at 30.0 kHz. Spikes were isolated online via manually-set thresholds for waveform detection triggers. As a measure of interhemispheric communication, we computed LFP-LFP coherence between left and right PRR over a broad range of frequencies. For each session using single-contact electrodes, we computed LFP-LFP coherence for each of the four possible interhemispheric pairings of electrodes. In each session using 32-channel multi-contact electrodes, we used LFP signals from every 4th contact (i.e., we used only 8 of the 32 contacts on each electrode, each separated by 800 μm) and we computed 4 average LFP-LFP coherence values from each of the 64 possible interhemispheric pairings of contacts.

### Quantification and statistical analysis

All data analyses were conducted using custom codes written in C, R, and MATLAB.

### LFP power

LFP power spectral density was estimated with a multitaper method. For each trial, the LFP signal was windowed with each of 4 orthogonal Slepian tapers and Fourier transforms were estimated. The Fourier transform of LFP signal *x_n_*(*t*) with the *k^th^* taper, *d_k_*(*t*) was estimated according to Eq. (1).

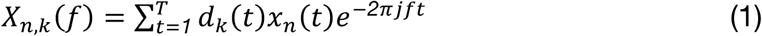

*x_n_*(*t*) is the windowed LFP signal for trial *n*, *T* is the length of *x_n_*(*t*), *f* is the frequency, and *j* is the imaginary unit 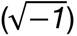. The power spectral density for a single trial *n*, *S_xx,n_* (*f*), was then estimated as a weighted average of auto-spectra |*X_n,k_*(*f*)|^2^ across tapers according to Eq. (2).

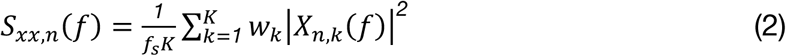

where *f_s_* is the sampling frequency, *K* is the number of tapers, and *w_k_* are weights determined by an adaptive algorithm (115). The power spectral density was then averaged across trials to produce a single estimate of the power spectral density according to Eq. (3) where *N* is the number of trials.

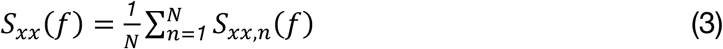

We used a time-half-bandwidth product of 2.5, affording us 4 Slepian tapers. We used either 400 ms or 200 ms windows, affording us frequency resolutions of ±6.25 or ±12.5 Hz, respectively. Band-limited power was estimated by summing the power spectral density estimate over the band of interest. Power time signals were estimated by stepping the time window by either 100 ms (400 ms windows) or 50 ms (200 ms windows) and estimating band-limited power at each time step. We present power time signals and power spectral density as percentage of baseline power or power spectral density, respectively. Baseline values are estimated as the average value over the 500 ms before target presentation. Power was computed at each LFP recording site individually before averaging across the population.

The bands of interest 20-30 Hz were not selected a priori. Instead, these bands were selected empirically early in our study to capture general trends in the power density spectra and then maintained as we collected more data. Note that with a frequency resolution of ±6.25 Hz (400 ms time windows), the band labeled 20-30 Hz includes information from frequencies from 14 Hz to 36 Hz. The same is true for measures of coherence described below.

### LFP-LFP coherence

We computed LFP-LFP coherence over a broad range of frequencies between left and right PRR during the last 500 ms of a variable delay period prior to the cue to initiate the reach.

Shared information between two LFP signals was quantified using coherence. Power spectral densities were estimated with the same multitaper method described above. Coherence between two LFP signals, *x* and *y*, was estimated according to Eq. (4).

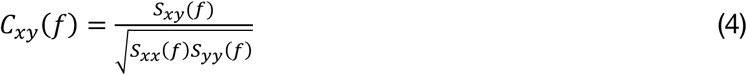

where *S_xx_*(*f*) and *S_yy_*(*f*) are the mean power density spectra across trials for LFP signals *x* and *y*, respectively, and *S_xy_*(*f*) is the cross spectrum for LFP signals *x* and *y*, averaged across all trials. The cross-spectrum for a single trial *n*, *S_xy,n_* (*f*) was estimated as a weighted average of the cross-spectra across tapers according to Eq. (5).

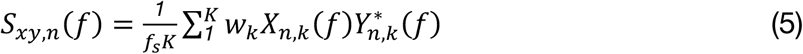

where *X_n,k_*(*f*) and *Y_n,k_*(*f*) are the Fourier transforms of time series *x*(*t*) and *y*(*t*), respectively, and 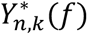 is the complex conjugate of *Y_n,k_*(*f*). The mean cross spectrum across trials is then estimated according to Eq. (6) where *N* is the number of trials.

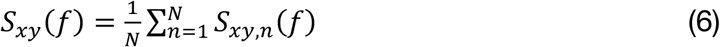

## Author Contributions

Experimental design: All authors. Investigation: J.K. and E.M. Formal analysis: J.K. and L.H.S. Original manuscript: J.K. Manuscript revision: All authors.

## Acknowledgements.

This work was supported by the National Eye Institute Grant EY-012135 (L.H.S.) and the Washington University Cognitive Computational and Systems Neuroscience Fellowship (J.K.).

## Data and Materials Sharing

Original data created for the study are or will be available in a persistent repository upon publication.

## Supplemental Information

**Fig. S1.**
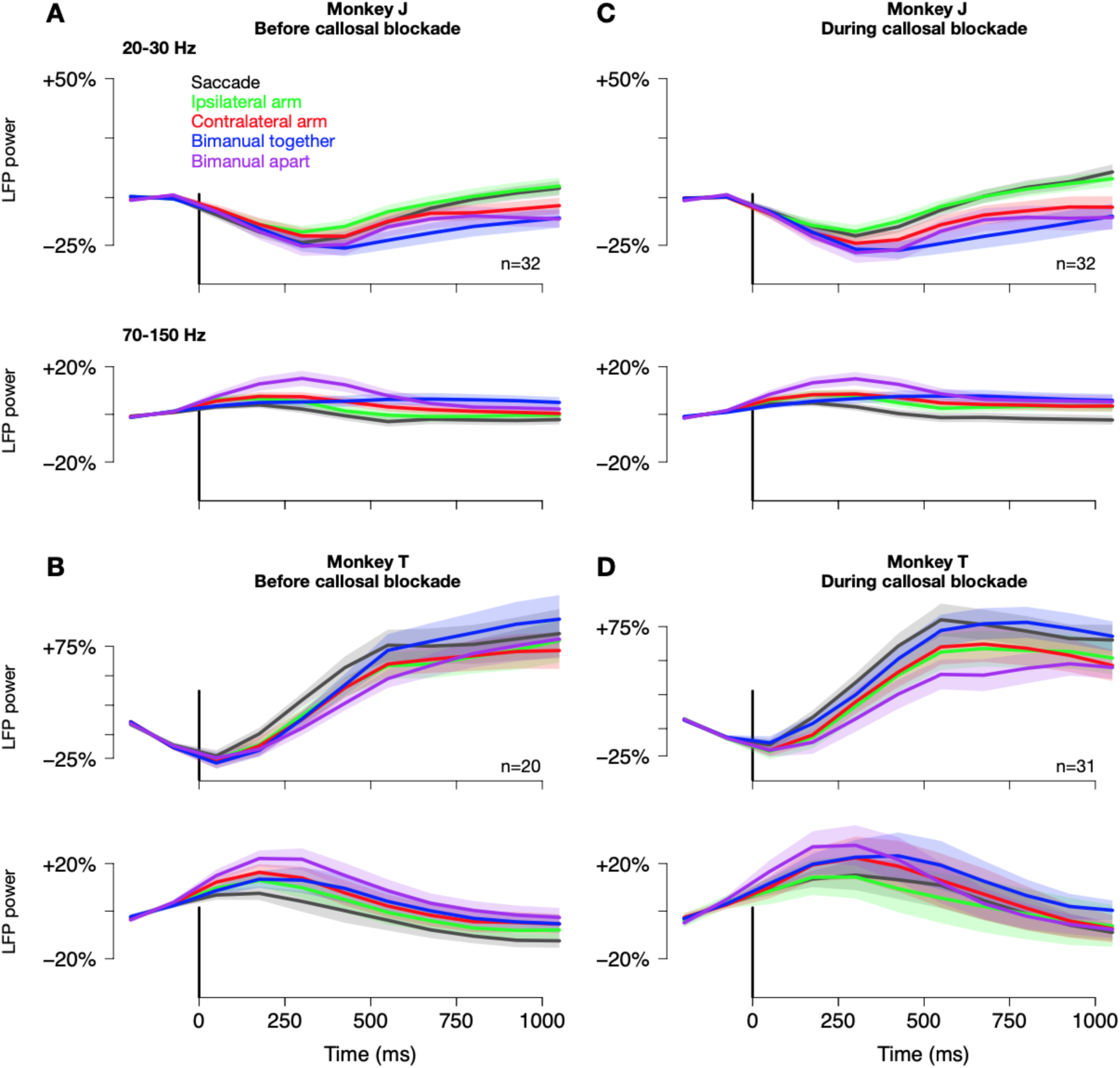
PRR LFP power aligned to target onset across the five movement tasks. *(A)* LFP power in 20-30 Hz (beta-band) and 70-150 Hz (gamma-band) in Monkey J before callosal blockade. *(B)* LFP power in 20-30 Hz and 70-150 Hz in Monkey J during callosal blockade. *(C)* LFP power in 20-30 Hz and 70-150 Hz in Monkey T before callosal blockade. *(D)* LFP power in 20-30 Hz and 70-150 Hz in Monkey T during callosal blockade. Shaded regions in all panels denote ±SEM. *n* denotes the number of sites for LFP recordings, which is much smaller than in the previous study where task-specific LFP power modulation (*n* = 312) was demonstrated (Mooshagian et al., 2021).

**Fig. S2.**
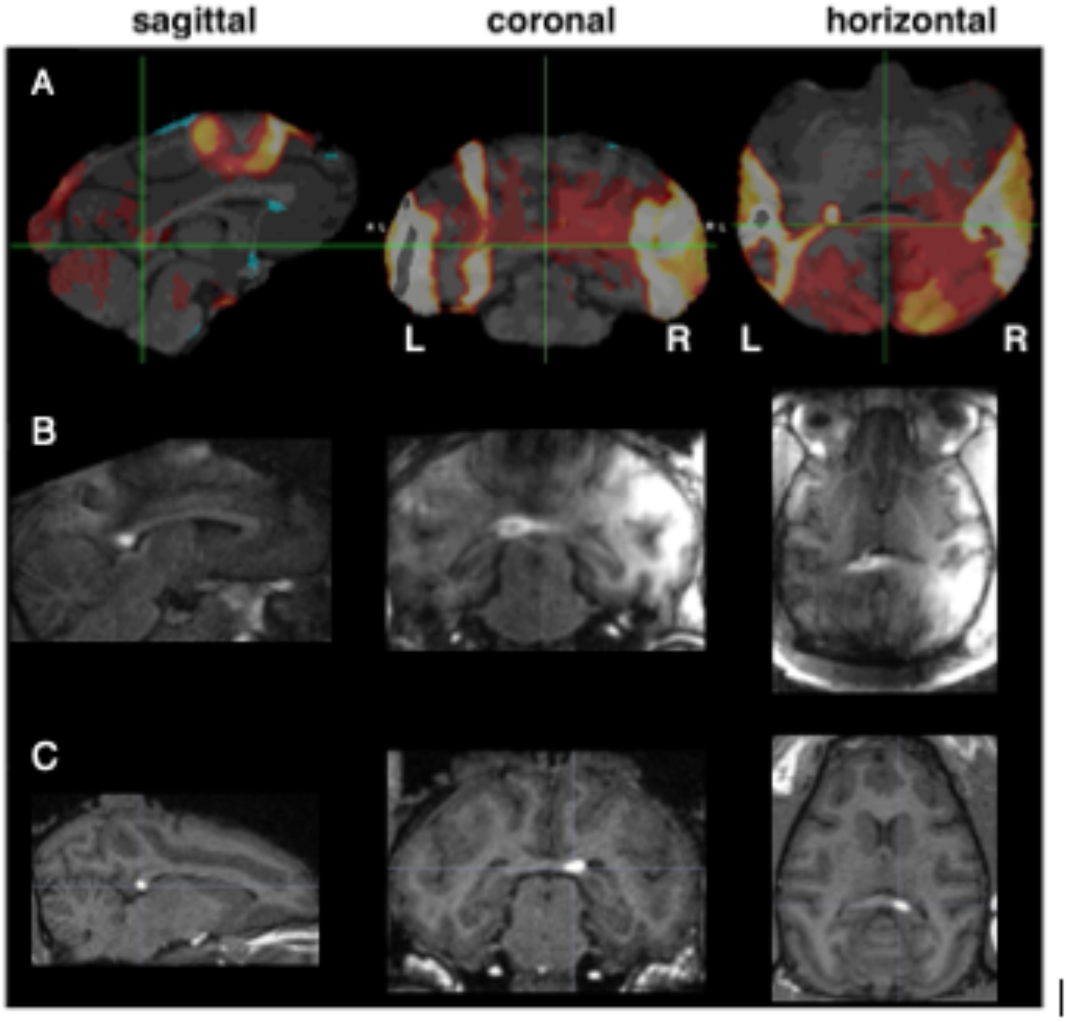
In vivo mapping of corpus callosum pathways using manganese-enhanced MRI. *(A)* Visualization of anterograde transport of manganese from right PRR in Monkey T. 1 μL of 0.1M MnCl_2_ buffered with 10 mM Tris–HCl was injected into right PRR. Structural MR images were acquired 24- and 48-hours post-injection. This is a subtraction MR image: 48 hr – 24 hr after the injection. The green crosshair indicates the callosal pathways from right PRR crossing to the left hemisphere. Callosal pathways connecting left and right PRR were restricted to the splenium. *(B)* Targeting of injections into the posterior callosum (Monkey T). Images are from awake scans acquired with a custom-made 8-channel coil. The image was acquired 6 hours after injecting 1 μL of 0.1M MnCl_2_ buffered with 10 mM Tris–HCl into the splenium. *(C)* Targeting of injections into the posterior callosum (Monkey J). Images are from an anesthetized scan acquired with 15-channel knee coil while the animal was anesthetized with isoflurane. The image was acquired 3 hours after injecting 0.08 μL of 0.1M MnCl_2_ buffered with 10 mM Tris–HCl into the splenium. We injected 2% lidocaine solution using the left recording chamber in Monkey T and the right chamber in Monkey J. We did not observe a lateralized effect; there was no behavioral effect specific to the contralateral arm to the side of the chamber used for 2% lidocaine injections.

**Fig. S3.**
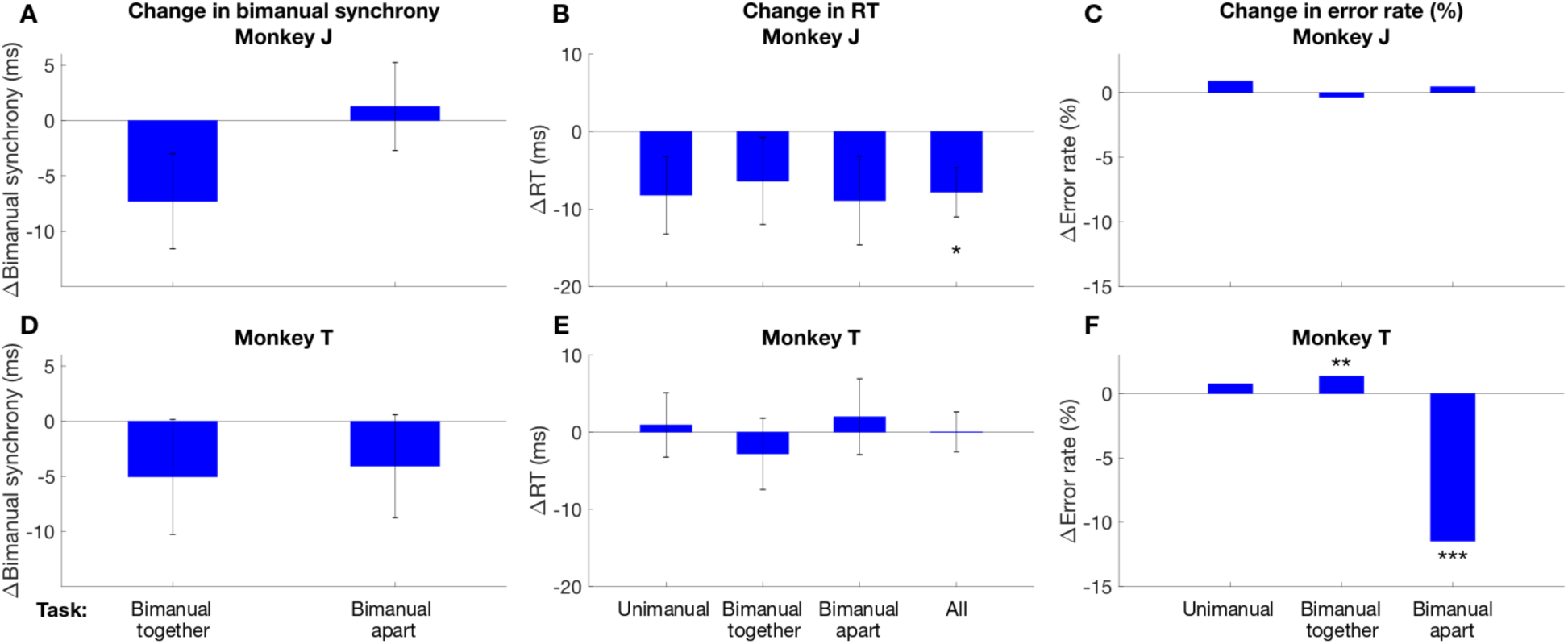
Behavioral performance in callosal blockade and control sessions in each animal. Format same as Fig. 3. *(A-C)* Monkey J. *(D-F)* Monkey T. *(A)* There was a trend in callosal blockade de-synchronizing bimanual-together movements in Monkey T by 7 ms (*p* = 0.1, pooled t-test). *(B)* Callosal blockade resulted in faster RT across all uni- and bimanual movements (*p* < 0.05, pooled t-test). *(C)* No significant change in error rate in Monkey J. *(D)* No significant change in bimanual synchrony in Monkey T. *(E)* No significant change in RT in Monkey T. *(F)* Callosal blockade worsened bimanual-together movements (*p* < 0.01, logistic regression) and improved bimanual-apart movements (*p* < 0.001, logistic regression) in Monkey T. Error bars indicate ±SEM. * *p* < 0.05; ** *p* < 0.01; *** *p* < 0.001

**Fig. S4.**
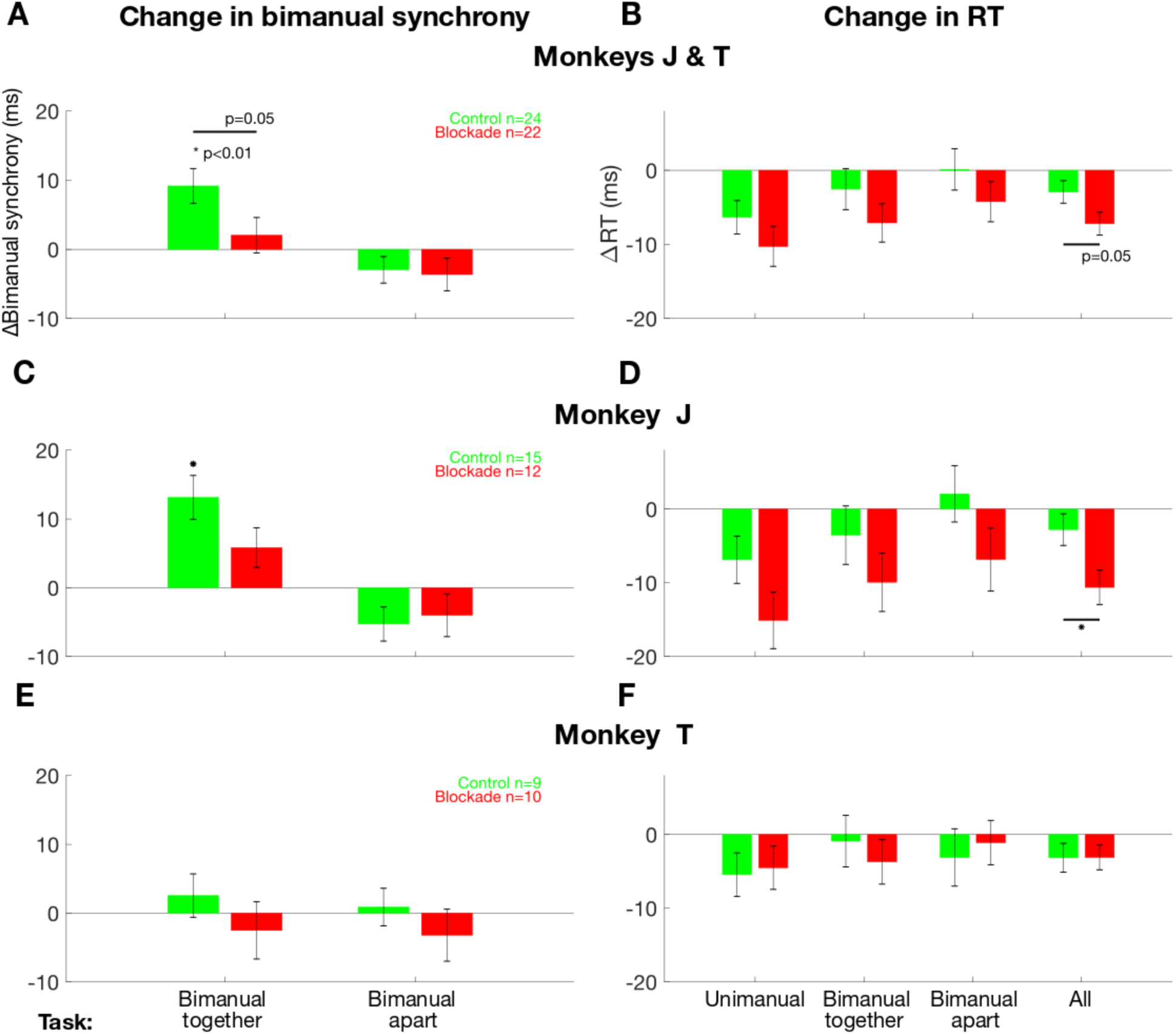
Bimanual synchrony and RT shown separately for control and blockade sessions. *(A)* Bimanual synchrony as a function of bimanual movement type and session in Monkeys J and T. We assayed the effect of callosal blockade on the absolute reaction time difference between the two arms, Δ(|RT_Left_ – RT_Right_|), in Peri minus Pre blocks. In control sessions, bimanual-together movements showed increased synchrony (decreased absolute timing difference) across blocks. This increase was significantly reduced in callosal blockade sessions (p = 0.05, two-sample t-test). No significant changes were observed in bimanual-apart trials. *(B)* Change in RT between blocks as a function of movement type and session in Monkeys J and T. Callosal blockade sped movement initiation (shorter RTs) compared to control for all three movement types. This speeding effect was statistically significant when averaged across all movement types (*p* = 0.05, two-sample t-test). ΔRT(Peri - Pre) is calculated as the difference between the median RT in the Peri block and the median RT in the Pre block for each movement type in both control and callosal blockade sessions. *(C)* Blockade desynchronized the two arms for bimanual together movements in Monkey J. In control sessions, the synchrony of bimanual together movements increased (the absolute timing difference decreased) in Peri compared to Pre blocks by −13 ms (paired t-test, *p* = 0.001). In callosal blockade sessions, there was no significant improvement in temporal synchrony of bimanual together movements, indicating a role of the callosum in bimanual movements to a single target. There was no effect on bimanual apart trials. *(D)* Blockade sped reach initiation in Monkey J. Blockade sped movements (shortens RTs) in all 3 cases compared to control, with a significant effect for All movement types (*p* < 0.02, two-sample t-test). *(E)* No significant difference in change in bimanual synchrony between control and callosal blockade sessions in Monkey T. *(F)* No significant difference in change in RT between control and callosal blockade sessions in Monkey T. Error bars in all panels denote ±SEM.

**Fig. S5.**
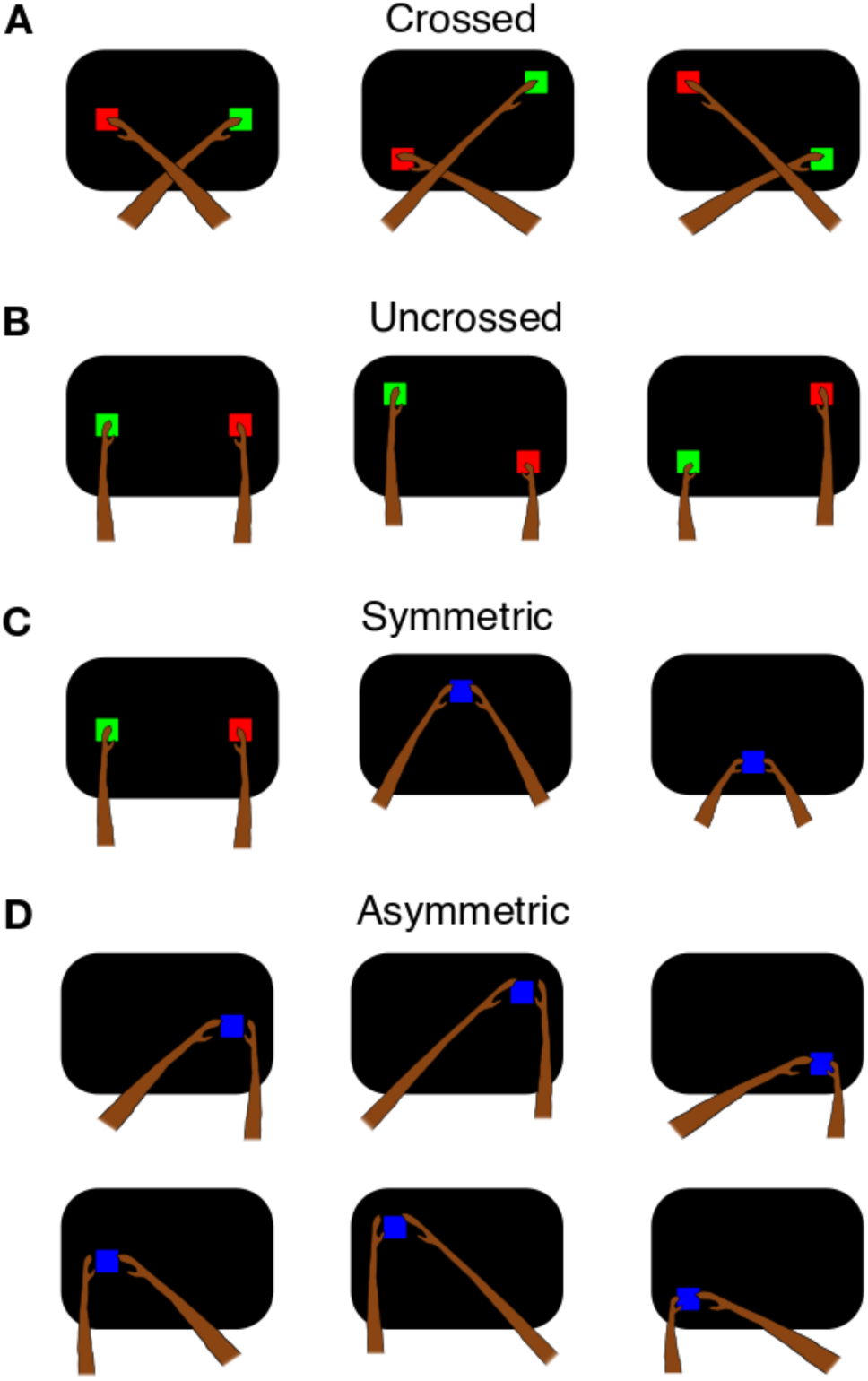
Bimanual movement types grouped by laterality (crossed or uncrossed) and symmetry (symmetric or asymmetric). *(A)* Crossed movements are bimanual-apart movements in which each arm reaches to a target in the contralateral visual hemifield (i.e., left hand to right visual field; right hand to left visual field). These movements require interhemispheric transfer because the hemisphere processing the spatial location of the target differs from the one generating the motor output. *(B)* Uncrossed movements are bimanual-apart movements in which each arm reaches to a target in the ipsilateral visual hemifield (i.e., left hand to left visual field; right hand to right visual field). Each hemisphere can process the spatial location of the target and generate the motor command locally, without requiring interhemispheric integration. *(C)* Symmetric movements are bimanual movements in which both arms move to spatially mirror-symmetric locations across the vertical midline. *(D)* Asymmetric movements are bimanual-together movements in which both arms move to the same lateralized target (e.g., both hands reach to a target in the left visual field), resulting in spatially asymmetric arm movements.

**Fig. S6.**
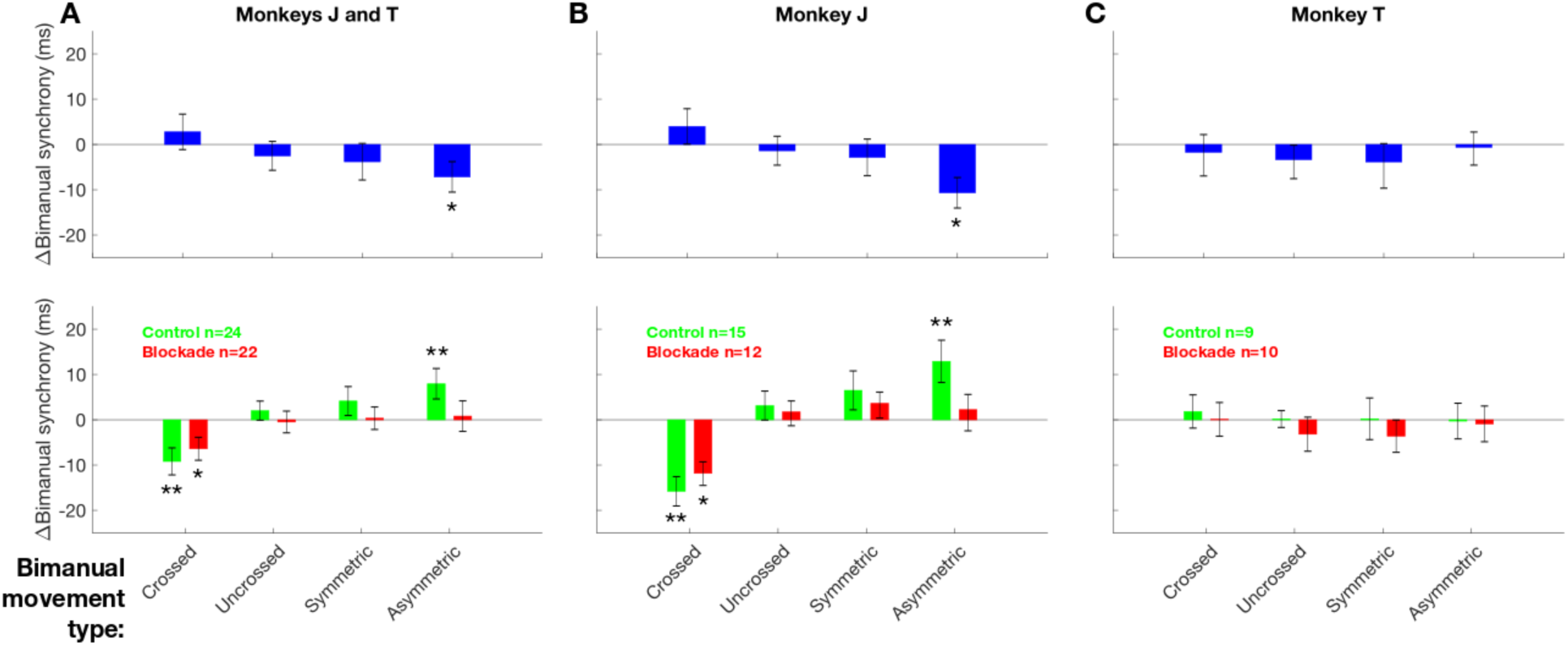
Mean change in movement synchrony across bimanual movement types grouped by laterality and symmetry. We quantified change in bimanual synchrony across different types of bimanual movements based on laterality and symmetry. Error bars in all panels denote ±SEM. Upper row: Format same as in Fig. 3A. We compared the change in bimanual synchrony from Pre to Peri blocks in control and blockade sessions. Statistics are based on two sample t-tests (* *p* < 0.05) Lower row: The change in bimanual synchrony from Pre to Peri blocks in control (green) and blockade (red) sessions shown separately. Statistics are based on paired t-tests (* *p* < 0.05; ** *p* < 0.01). *(A)* In Monkeys J and T, bimanual synchrony for asymmetric bimanual movements was significantly worse in blockade sessions compared to control sessions (blue bar; −7.2±3.4 ms; *p* < 0.05). Crossed movements were less synchronous in Peri blocks in both control (*p* < 0.01) and blockade (*p* < 0.05) sessions. Asymmetric movements were more synchronous in Peri blocks in control sessions (*p* < 0.01). *(B)* In Monkey J, bimanual synchrony for asymmetric bimanual movements was significantly worse in blockade sessions compared to control sessions (blue bar; −10.7±4.7 ms; *p* < 0.05). Crossed movements were less synchronous in Peri blocks in both control (*p* < 0.01) and blockade (*p* < 0.05) sessions. Asymmetric movements were more synchronous in Peri blocks in control sessions (*p* < 0.01). *(C)* No significant effect in bimanual synchrony in Monkey T.

**Fig. S7.**
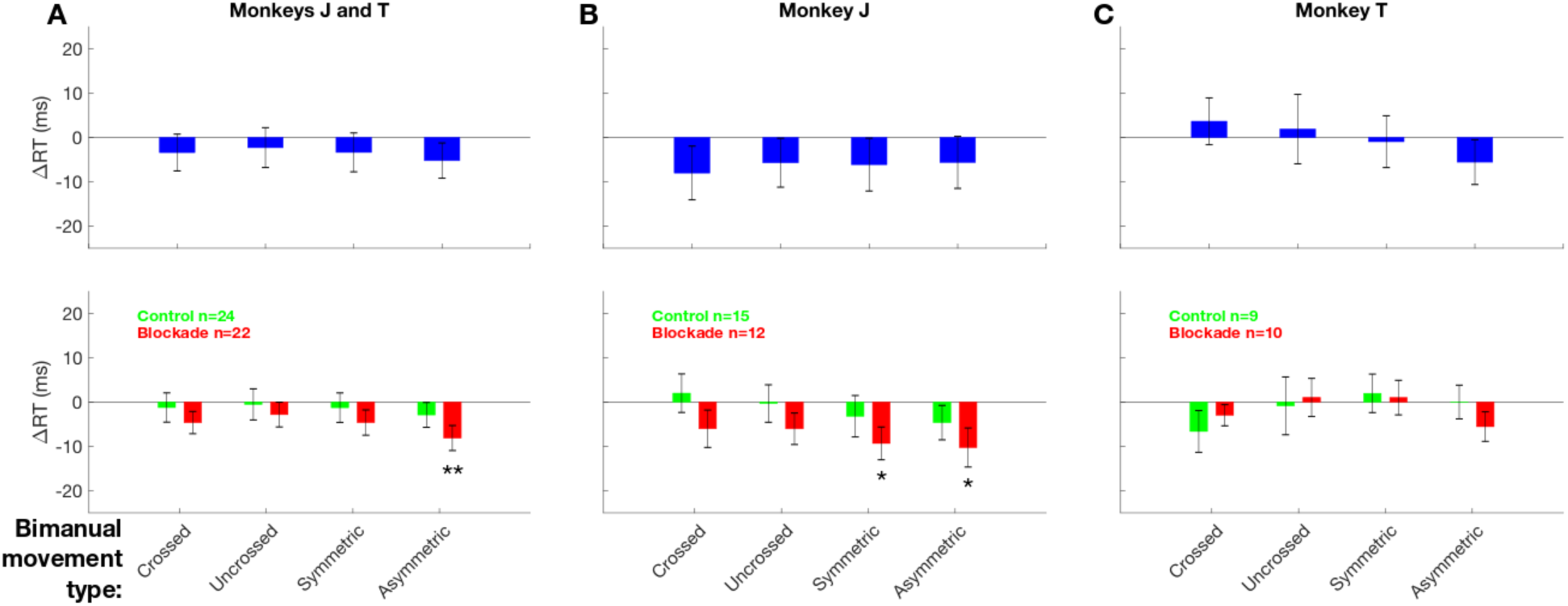
Change in RT in bimanual movement types grouped by laterality and symmetry. We quantified change in RT across different types of bimanual movements as grouped in Fig. S6. Error bars in all panels denote ±SEM. Upper row: Format same as in Fig. 3B. We compared the change in RT from Pre to Peri blocks in control and blockade sessions. Lower row: The change in RT from Pre to Peri blocks in control (green) and blockade (red) sessions shown separately. Statistics are based on paired t-tests (** *p* < 0.01; * *p* < 0.05). *(A)* In Monkeys J and T, there was no significant change in RT when comparing blockade vs. control sessions (blue bars) or in blockade (red) and control (green) sessions separately except for asymmetric movements in blockade sessions (*p* < 0.01). *(B)* In Monkey J, there was no significant change in RT when comparing blockade and control sessions (blue bars). However, symmetric and asymmetric movements had faster RT in Peri blocks in blockade sessions (*p* < 0.05). *(C)* No significant effect in RT in Monkey T.

**Fig. S8.**
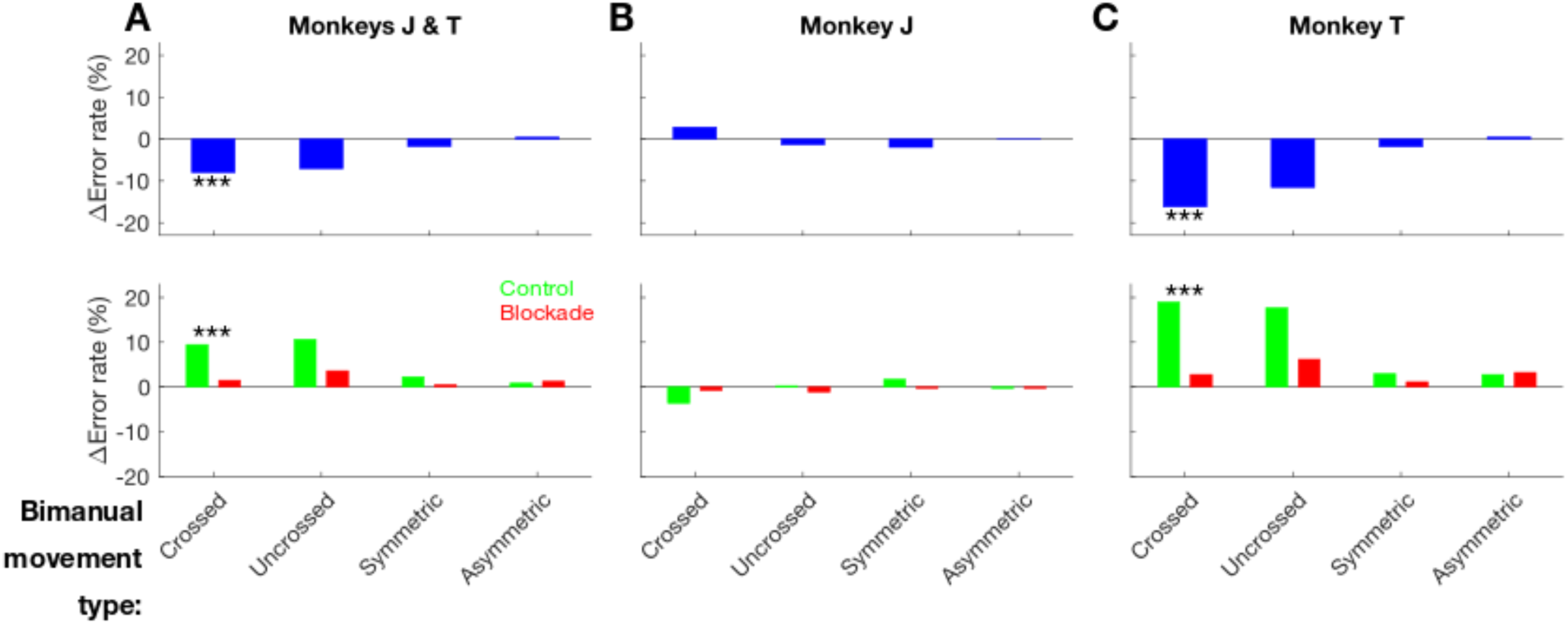
Change in arm error rates in bimanual movement types grouped by laterality and symmetry. We quantified change in error rates across different types of bimanual movements as grouped in Fig. S6. ΔError rate (Peri - Pre) is calculated as the difference between error rates in the Peri block and Pre block. Upper row: We compared the change in error rates in control and blockade sessions. Lower row: ΔError rate (Peri - Pre) shown separately for control and blockade sessions. *(A)* Crossed arm movements significantly improved by 8% (*p* < 0.001, logistic regression) in callosal blockade sessions in Monkeys J and T. *(B)* No clear effect on error rate in Monkey J. *(C)* Crossed arm movements significantly improved by 16% (*p* < 0.001, logistic regression) in callosal blockade sessions in Monkey T.

**Fig. S9.**
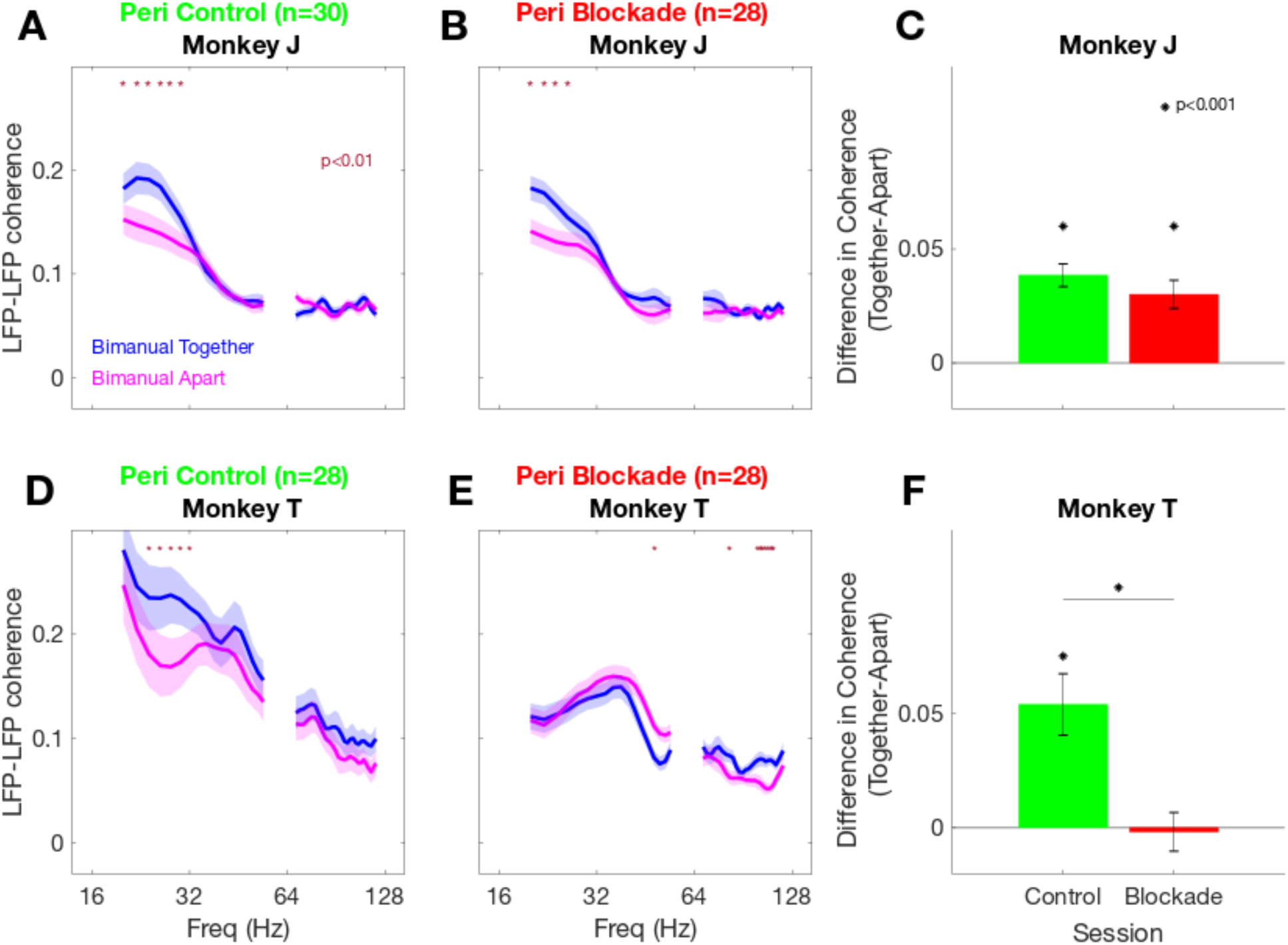
LFP-LFP coherence in each animal. Format same as in Fig. 4. *(A-C)* Monkey J. *(D-F)* Monkey T. *(A)* During Peri control blocks, coherence is higher for bimanual-together movements than for bimanual-apart movements over the beta frequency range (20 – 30 Hz). *(B)* During Peri callosal blockade blocks, the bimanual-task specific difference in coherence in 20 to 30 Hz is reduced. *(C)* Blocking the corpus callosum reduced the task-dependent difference in coherence, but the effect was not significant (*p* = 0.29, pooled t-test). *(D)* During Peri control blocks, LFP-LFP coherence in bimanual together movements has a higher coherence than bimanual apart movements in 20-30 Hz. *(E)* During Peri callosal blockade blocks, the difference in coherence between the two bimanual movement tasks was significantly reduced in 20-30 Hz. *(F)* Difference in coherence (Together-Apart) in 20 to 30 Hz are statistically significant in Peri control (*p* < 0.001, paired t-test) and but not in Peri callosal blockade sessions. The difference in coherence (Together-Apart) is significantly reduced in callosal blockade sessions (red) compared to control sessions (green) (*p* < 0.001, pooled t-test).

**Fig. S10.**
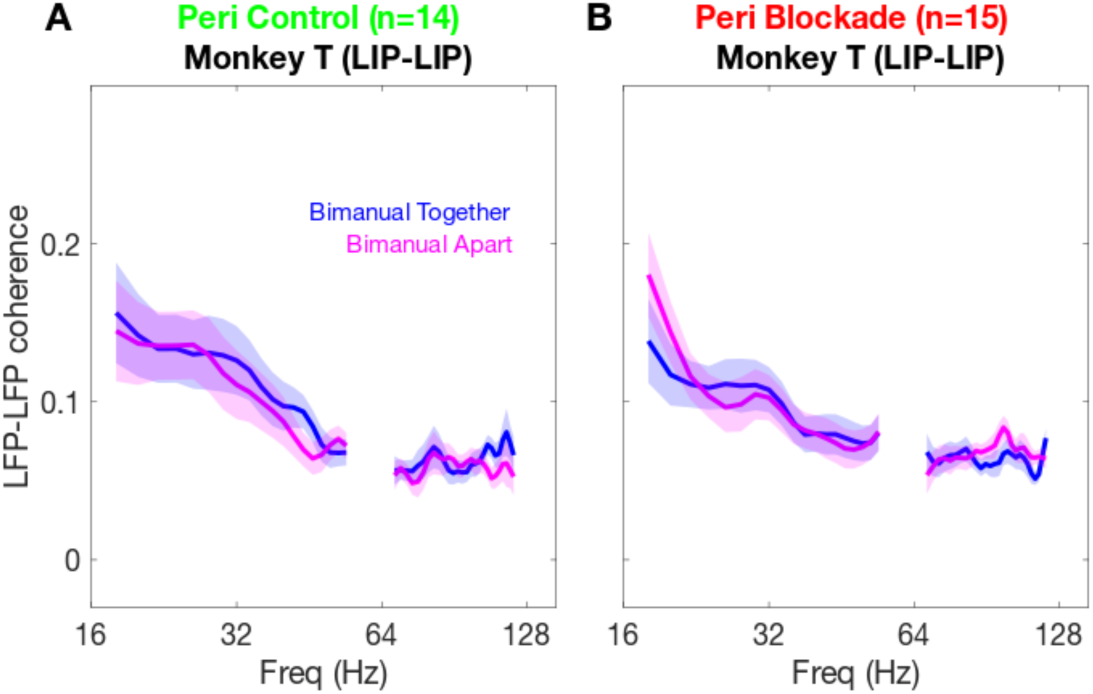
LFP-LFP coherence between left and right LIP in a delayed memory task (Monkey T). Format same as in Fig. 4A–B. There was no significant task-specific modulation in LFP-LFP coherence in 20-30 Hz between left and right LIP in (A) Peri Control (0.014±0.012) and (B) Peri Blockade (0.003±0.009) in Monkey T.

**Table S1.** Mean eye and arm error rates (%). *(A)* Most eye errors occurred before the Go Cue and were due to premature saccade initiation. *(B)* Eye errors after the Go Cue were due to inaccurate saccade endpoints. *(C)* Arm errors before the Go Cue were due to premature arm movement initiation. *(D)* Arm movement errors after the Go Cue were due to inaccurate reach endpoints. Most arm errors occurred after Go Cue. Error rates are lower than those shown in Fig. 3 because the error rates in this table are averaged over sessions while the error rates in Fig. 3 are calculated by merging all success and error trials into a single session for binomial test. Because of the scarcity of arm movement errors, we used the binomial test in the main analyses.

| Error type | Session |  |  |  |
| --- | --- | --- | --- | --- |
| | Control ($n = 24$) | | Blockade ($n = 22$) | |
| Movement type | Pre | Peri | Pre | Peri |
| **A** Eye movement, before Go Cue: |  |  |  |  |
| Saccade-only | 14 | 17 | 15 | 16 |
| Unimanual | 15 | 14 | 12 | 11 |
| Bimanual-together | 16 | 15 | 13 | 15 |
| Bimanual-apart | 11 | 12 | 11 | 11 |
| **B** Eye movement, after Go Cue: |  |  |  |  |
| Saccade-only | 2 | 2 | 2 | 3 |
| Unimanual | 1 | 1 | 1 | 1 |
| Bimanual-together | 1 | 1 | 1 | 0 |
| Bimanual-apart | 0 | 0 | 0 | 0 |
| **C** Arm movement, before Go Cue: |  |  |  |  |
| Saccade-only | 0 | 1 | 0 | 0 |
| Unimanual | 0 | 1 | 1 | 1 |
| Bimanual-together | 0 | 1 | 1 | 1 |
| Bimanual-apart | 0 | 1 | 1 | 0 |
| **D** Arm movements, after Go Cue: |  |  |  |  |
| Saccade-only | 1 | 1 | 0 | 0 |
| Unimanual | 3 | 3 | 2 | 3 |
| Bimanual-together | 4 | 4 | 3 | 5 |
| Bimanual-apart | 4 | 8 | 3 | 5 |

Values are shown as the mean percentage.

